# The lectin-specific activity of *Toxoplasma gondii* microneme proteins 1 and 4 binds Toll-like receptor 2 and 4 N-glycans to regulate innate immune priming

**DOI:** 10.1101/187690

**Authors:** Aline Sardinha-Silva, Flávia C. Mendonça-Natividade, Camila F. Pinzan, Carla D. Lopes, Diego L. Costa, Damien Jacot, Fabricio F. Fernandes, André L. V. Zorzetto-Fernandes, Nicholas J. Gay, Alan Sher, Dragana Jankovic, Dominique Soldati-Favre, Michael E. Grigg, Maria Cristina Roque-Barreira

**Affiliations:** Department of Cell and Molecular Biology and Pathogenic Bioagents, Ribeirão Preto Medical School, University of São Paulo-USP (FMRP/USP), Ribeirão Preto, São Paulo, 14049-900, Brazil; Laboratory of Parasitic Diseases, National Institute of Allergy and Infectious Diseases, National Institutes of Health, Bethesda, MD 20892, USA; Department of Microbiology and Molecular Medicine, CMU, University of Geneva, 1 rue Michel-Servet, 1211 Geneva 4, Switzerland; Department of Biochemistry, Cambridge University, 80 Tennis Court Road Cambridge CB2 1GA, United Kingdom

## Abstract

Infection of host cells by *Toxoplasma gondii* is an active process, which is regulated by secretion of microneme (MICs) and rhoptry proteins (ROPs and RONs) from specialized organelles in the apical pole of the parasite. MIC1, MIC4 and MIC6 assemble into an adhesin complex, secreted on the parasite surface and function to promote infection competency. MIC1 and MIC4 are known to bind terminal sialic acid residues and galactose residues, respectively and to induce IL-12 production from splenocytes. Here we show that rMIC1- and rMIC4-stimulated dendritic cells and macrophages to produce proinflammatory cytokines, and they do so by engaging TLR2 and TLR4. This process depends on sugar recognition, since point mutations in the carbohydrate-recognition domains (CRD) of rMIC1 and rMIC4 inhibit innate immune cells activation. HEK cells transfected with TLR2 glycomutants were selectively unresponsive to MICs. Following *in vitro* infection, parasites lacking MIC1 or MIC4, as well as expressing MIC proteins with point mutations in their CRD, failed to induce wild-type (WT) levels of IL-12 secretion by innate immune cells. However, only MIC1 was shown to impact systemic levels of IL-12 and IFN-γ *in vivo*. Together, our data show that MIC1 and MIC4 interact physically with TLR2 and TLR4 N-glycans to trigger IL-12 responses, and MIC1 is playing a significant role *in vivo* by altering *T. gondii* infection competency and murine pathogenesis.

**AUTHOR SUMMARY:** Toxoplasmosis is caused by the protozoan *Toxoplasma gondii*, belonging to the Apicomplexa phylum. This phylum comprises important parasites able to infect a broad diversity of animals, including humans. A particularity of *T. gondii* is its ability to invade virtually any nucleated cell of all warm-blooded animals through an active process, which depends on the secretion of adhesin proteins. These proteins are discharged by specialized organelles localized in the parasite apical region, and termed micronemes and rhoptries. We show in this study that two microneme proteins from *T. gondii* utilize their adhesion activity to stimulate innate immunity. These microneme proteins, denoted MIC1 and MIC4, recognize specific sugars on receptors expressed on the surface of mammalian immune cells. This binding activates these innate immune cells to secrete cytokines, which promotes efficient host defense mechanisms against the parasite and regulate their pathogenesis. This activity promotes a chronic infection by controlling parasite replication during acute infection.

## INTRODUCTION

*Toxoplasma gondii* is a coccidian parasite belonging to the phylum Apicomplexa and is the causative agent of toxoplasmosis. This protozoan parasite infects a variety of vertebrate hosts, including humans with about one-third of the global population being chronically infected [1]. Toxoplasmosis can be fatal in immunocompromised individuals or when contracted congenitally [1], and is considered the second leading cause of death from foodborne illnesses in the United States [2].

*T. gondii* invades host cells through an active process that relies on the parasite actinomyosin system, concomitantly with the release of microneme proteins (MICs) and rhoptry neck proteins (RONs) from specialized organelles in the apical pole of the parasite [3]. These proteins are secreted by tachyzoites [4, 5] and form complexes composed of soluble and transmembrane proteins. Some of the MICs act as adhesins, interacting tightly with host cell-membrane glycoproteins and receptors, and are involved in the formation of the moving junction [6]. This sequence of events ensures tachyzoite gliding motility, migration through host cells, invasion and egress from infected cells [4, 7]. Among the released proteins, MIC1, MIC4, and MIC6 form a complex that, together with other *T. gondii* proteins, plays a role in the adhesion and invasion of host cells [8, 9], contributing to the virulence of the parasite [10, 11].

Several studies have shown that host-cell invasion by apicomplexan parasites such as *T. gondii* involves carbohydrate recognition [12–15]. Interestingly, MIC1 and MIC4 have lectin domains [11, 16–18] that recognize oligosaccharides with sialic acid and D-galactose in the terminal position, respectively. Importantly, the parasite’s Lac^+^ subcomplex, consisting of MIC1 and MIC4, induces adherent spleen cells to release IL-12 [17], a cytokine critical for the protective response of the host to *T. gondii* infection [19]. In addition, immunization with this native subcomplex, or with recombinant MIC1 (rMIC1) and MIC4 (rMIC4), protects mice against experimental toxoplasmosis [20, 21]. The induction of IL-12 is typically due to detection of the pathogen by innate immunity receptors, including members of the Toll-like receptor (TLR) family, whose stimulation involves MyD88 activation and priming of Th1 responses, which protects the host against *T. gondii* [19, 22]. It is also known that dysregulated expression of IL-12 and IFN-γ during acute toxoplasmosis can drive a lethal immune response, in which mice succumb to infection by severe immunopathology, the result of insufficient levels of IL-10 and/or a collapse in the regulatory CD4+Foxp3+ T cell population [23, 24].

Interestingly, regarding the innate immune receptors associated with IL-12 response during several infections, the extracellular leucine-rich repeat domains of TLR2 and TLR4 contain four and nine N-glycans, respectively [25]. Therefore, we hypothesized that MIC1 and MIC4 bind TLR2 and TLR4 N-glycans on antigen-presenting cells (APCs) and, through this interaction, trigger immune cell activation and IL-12 production. To investigate this possibility, we assayed the ability of rMIC1 and rMIC4 to bind and activate TLR2 and TLR4. Using several strategies, we demonstrated that TLR2 and TLR4 are indeed critical targets for both MIC1 and MIC4. These parasite and host cell structures establish lectin-carbohydrate interactions that contribute to the induction of IL-12 production by innate immune cells, and we show here that the MIC1 lectin promotes *T. gondii* infection competency and regulates parasite virulence during *in vivo* infection.

## RESULTS

### Lectin properties of recombinant MIC1 and MIC4 are consistent with those of the native Lac^+^ subcomplex

The native MIC1/4 subcomplex purified from soluble *T. gondii* antigens has lectin properties, so we investigated whether their recombinant counterparts retained the sugar-binding specificity. The glycoarray analysis revealed the interactions of: i) the Lac^+^ subcomplex with glycans containing terminal α(2-3)-sialyl and β(1-4)- or β(1-3)-galactose; ii), rMIC1 with α(2-3)-sialyl residues linked to β-galactosides; and iii) of rMIC4 with oligosaccharides with terminal β(1-4)- or β(1-3)-galactose (Fig 1A). The combined specificities of the individual recombinant proteins correspond to the dual sugar specificity of the Lac^+^ fraction, demonstrating that the sugar-recognition properties of the recombinant proteins are consistent with those of the native ones.

**Fig 1.**
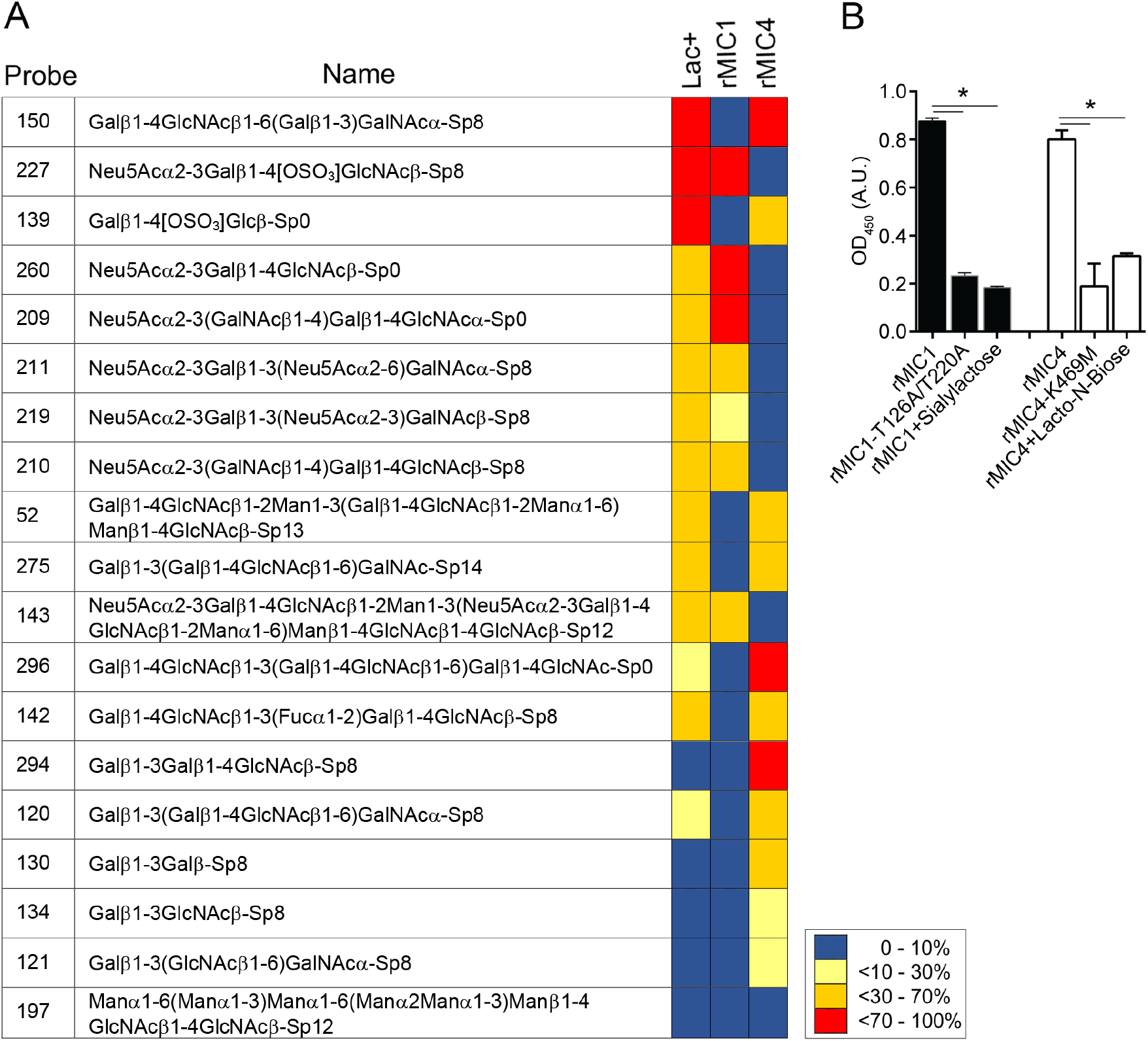
Lectin activity of rMIC1 and rMIC4. **(A)** Glycoarray of the native MIC1/MIC4 subcomplex (Lac+) and of the recombinant forms of MIC1 and MIC4. In total, 320 oligosaccharide probes were analysed by reading their fluorescence intensities, and the 20 best recognized glycans are shown. The results were represented as previously reported [18]. **(B)** The activity and inhibition assays were performed in microplates coated with glycoproteins with or without sialic acid, fetuin (black bars), or asialofetuin (white bars), separately. After coating, wild type or mutated rMIC1 and rMIC4, pre-incubated with PBS or their corresponding sugars, were added to the wells. Later, bound proteins were detected through the addition of serum from *T. gondii-*infected mice. Data in **(B)** are expressed as mean ±S.D. of triplicate wells and significance was calculated with ANOVA followed by Bonferroni’s multiple comparisons test. *p<0.05. Data are representative of two **(B)** independent experiments. Gal: galactose; GalNAc: *N-* acetylgalactosamine; Glc: glucose; Man: mannose; Fuc: fucose; Neu5Ac: *N-* acetylneuraminic acid; wt: wild type protein; mut: protein with a mutation in the carbohydrate-recognition domain (CRD); ns: not significant.

Based on the sugar recognition selectivity of rMIC1 and rMIC4, we tested two oligosaccharides (α(2-3)-sialyllactose and lacto-N-biose) for their ability to inhibit the interaction of the MICs with the glycoproteins fetuin and asialofetuin [26]. Sialyllactose inhibited the binding of rMIC1 to fetuin, and lacto-N-biose inhibited the binding of rMIC4 to asialofetuin (Fig 1B). To ratify the carbohydrate recognition activity of rMIC1 and rMIC4, we generated point mutations into the carbohydrate recognition domains (CRDs) of the rMICs to abolish their lectin properties [11, 18, 27]. These mutated forms, i.e. rMIC1-T126A/T220A and rMIC4-K469M, lost their capacity to bind to fetuin and asialofetuin, respectively (Fig 1B), having absorbance as low as that in the presence of the specific sugars. Thus, our results indicate that rMIC1 and rMIC4 maintained their lectin properties, and that the CRD function can be blocked either by competition with specific sugars or by targeted mutations.

### rMIC1 and rMIC4 trigger the activation of DCs and macrophages

We have previously demonstrated that the native Lac^+^ subcomplex stimulates murine adherent spleen cells to produce proinflammatory cytokines [20]. We evaluated whether recombinant MIC1 and MIC4 retained this property and exerted it on BMDCs and BMDMs. BMDCs (Fig 2A-2D) and BMDMs (Fig 2E-2H) produced high levels of the proinflammatory cytokines IL-12 (Fig 2A and 2E), TNF-α (Fig 2B and 2F), and IL-6 (Fig 2C and 2G). This was not attributable to residual LPS contamination as the recombinant protein assays were done in the presence of polymyxin B, and LPS levels were less than 0.5ng/ml [see Materials and Methods section]. Although conventional CD4^+^ Th1 cells are known to be the major producers of IL-10 during murine *T. gondii* infection [28], we also found that rMIC1 and rMIC4 induced the production of this cytokine by BMDCs (Fig 2D) and BMDMs (Fig 2H). We verified that the two recombinant proteins induced the production of similar levels of IL-12, TNF-α, and IL-6 by both BMDCs (Fig 2A-2C) and BMDMs (Fig 2E-2G). Both MICs induced the production of similar levels of IL-10 in BMDCs (Fig 2D); however, BMDMs produced significantly higher levels of IL-10 when stimulated with rMIC1 than when stimulated with rMIC4 (Fig 2H). These cytokine levels were similar to those induced by the TLR4 agonist LPS. Thus, recombinant MIC1 and MIC4 induce a proinflammatory response in innate immune cells, which is consistent with the results obtained for the native Lac^+^ subcomplex [20].

**Fig 2.**
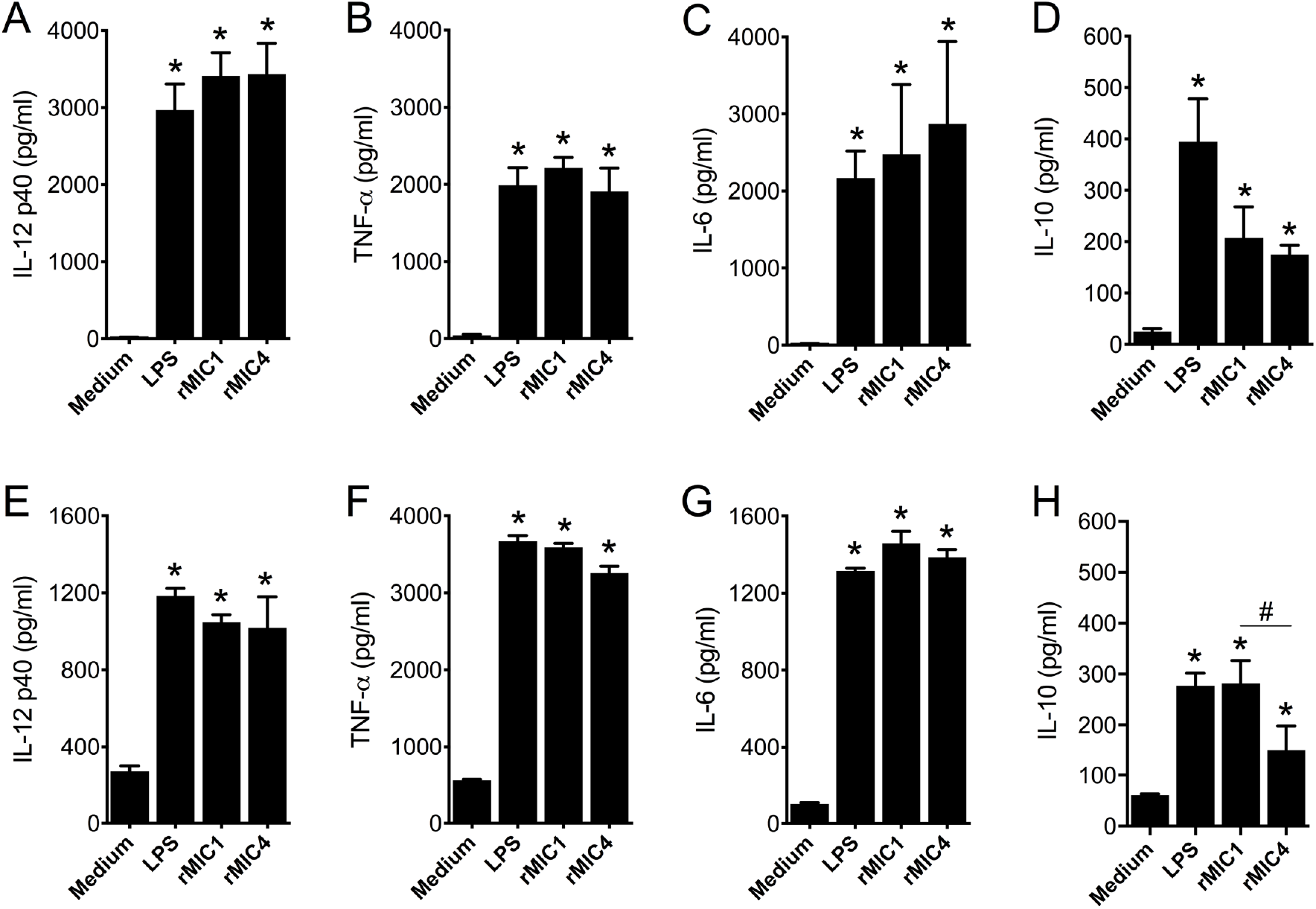
Microneme proteins stimulate cytokine production by dendritic cells and macrophages. **(A-D)** Bone marrow-derived dendritic cells (BMDCs) and **(E-H)** bone marrow-derived macrophages (BMDMs) from C57BL/6 mice were stimulated with rMIC1 (5 μg/mL) and rMIC4 (5 μg/mL) for 48 h. LPS (100 ng/mL) was used as positive control. The levels of IL-12p40, TNF-α, and IL-6 were measured by ELISA. For this assay, rMIC1 and rMIC4 were passed through polymyxin B column, followed by incubation with polymyxin B sulphate salt media preparation that was added to the culture (see Material and Methods). Data are expressed as mean ±S.D. of triplicate wells and significance was calculated with ANOVA followed by Bonferroni’s multiple comparisons test. *p<0.05. Data are representative of three independent experiments.

### The activation of macrophages by rMIC1 and rMIC4 depends on TLR2 and TLR4

To investigate the mechanisms through which *T. gondii* MIC1 and MIC4 stimulate innate immune cells to produce cytokines, we assessed whether these MICs can activate specific TLRs. To this end, BMDMs from WT, MyD88^−/−^, TRIF^−/−^, TLR2^−/−^, TLR4^−/−^, or TLR2/4 DKO mice, as well as HEK293T cells transfected with TLR2 or TLR4, were cultured in the presence or absence of rMIC1 and rMIC4 for 48 hours. The production of IL-12 by BMDMs (Fig 3A-3I) and IL-8 by HEK cells (Fig 3J and 3K) were used as an indicator of cell activation. IL-12 production by BMDMs from MyD88^−/−^, TRIF^−/−^, TLR2^−/−^, and TLR4^−/−^ mice was lower than that of BMDMs from WT mice (Fig 3A-3D); no IL-12 was detected in cultures of TLR2/4 DKO mice cells stimulated with either rMIC1 or rMIC4 (Fig 3E). These results show that TLR2 and TLR4 are both relevant for the activation of macrophages induced by rMIC1 and rMIC4. The residual cytokine production observed in macrophages from TLR2^−/−^ or MyD88^−/−^ mice may be the result of activation of TLR4 (Fig 3A and 3C), and vice versa; e.g., the residual IL-12 levels produced by macrophages from TLR4^−/−^ mice may be the result of TLR2 activation. The finding that MICs fail to induce IL-12 production in DKO mice BMDMs suggests that cell activation triggered by *T. gondii* MIC1 or MIC4 does not require the participation of other innate immunity receptors beyond TLR2 and TLR4. Nevertheless, because parasite components such as DNA or profilin engage TLR9, TLR11, and TLR12 to produce IL-12 in macrophages [19, 22, 29], we investigated the involvement of these receptors, as well as TLR3 and TLR5, in the response to rMIC1 or rMIC4. BMDMs from TLR3^−/−^, TLR5^−/−^, TLR9^−/−^, and TLR11/12 DKO mice stimulated with rMIC1 or rMIC4 produced similar levels of IL-12 as cells from WT (Fig 3F-3I), indicating that the activation triggered by rMIC1 or rMIC4 does not depend on these receptors. Additionally, stimulation of HEK cells transfected with human TLR2 (Fig 3J) or TLR4 (Fig 3K) with optimal concentrations of rMIC1 (Fig S1A and S1C) and rMIC4 (Fig S1B and S1D) induced IL-8 production at levels that were higher than those detected in the absence of stimuli (medium), and similar to those induced by the positive controls. Finally, by means of a pull-down experiment, we demonstrated a physical interaction between rMIC1 and TLR2 or TLR4 and between rMIC4 and TLR2 or TLR4 (Fig 3L).

**Fig 3.**
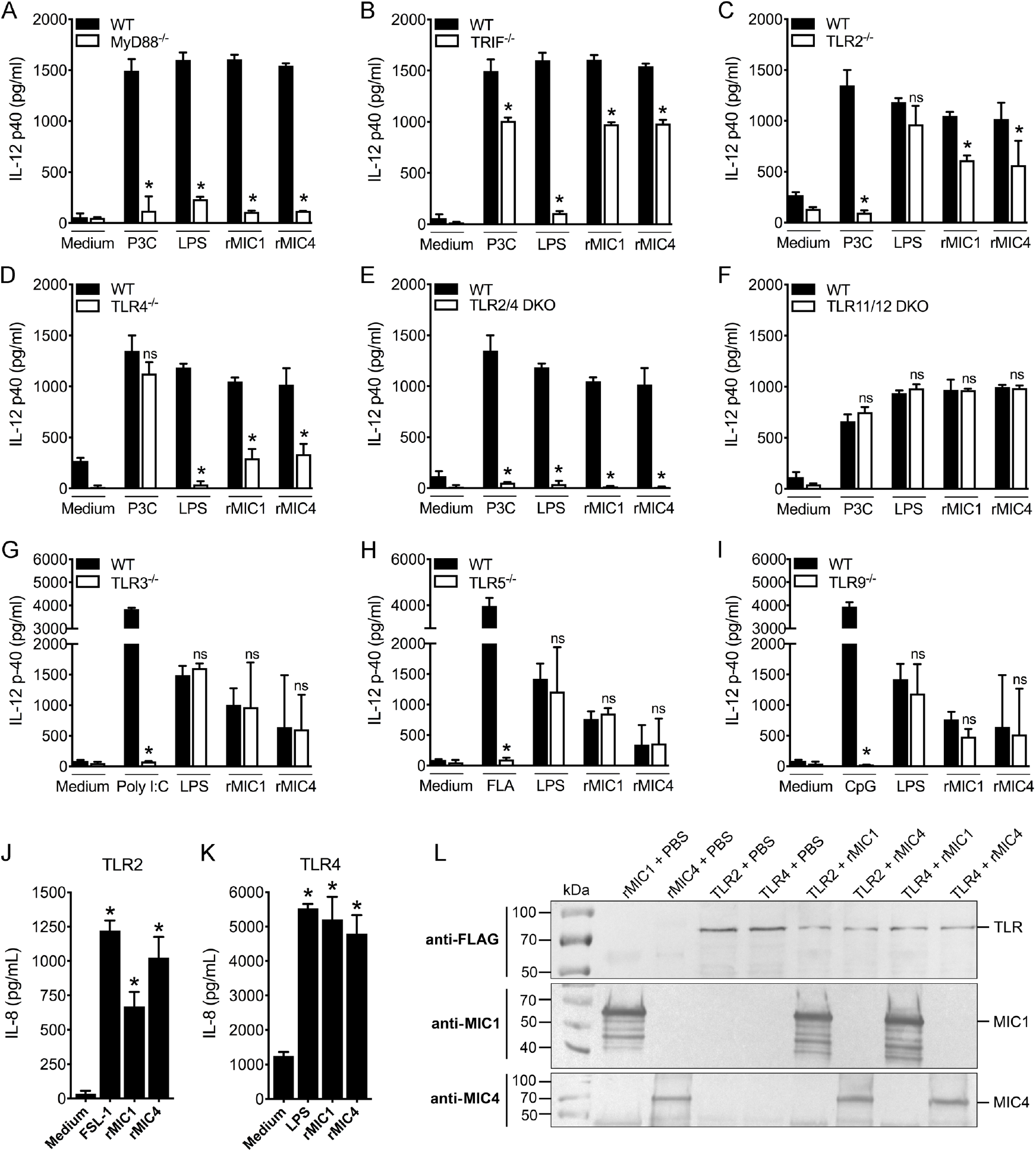
The IL-12 production induced by rMICs is dependent on binding to TLR2 and TLR4. **(A-I)** Bone marrow-derived macrophages from WT, TLR2^−/−^, TLR4^−/−^, double knockout TLR2^−/−^/TLR4^−/−^, TLR3^−/−^, TLR5^−/−^, TLR9^−/−^, and double knockout TLR11^−/−^/TLR12^−/−^ mice, all of the C57BL/6 background, were stimulated with rMIC1 or rMIC4 (5 μg/mL) for 48 h. Pam3CSK4 (P3C) (1 μg/mL), LPS (100 ng/mL), Poly I:C (10 μg/mL), Flagellin (FLA) (1 μg/mL) and CpG (25 μg/mL) were used as positive controls. IL-12p40 levels were measured by ELISA. **(J and K)** Transfected HEK293T cells expressing TLR2 were stimulated with rMIC1 (750 nM) or rMIC4 (500 nM), and rMIC1 (200 nM) or rMIC4 (160 nM) for HEK cells expressing TLR4, for 24 h. FSL-1 (100 ng/mL) and LPS (100 ng/mL) were used as positive controls. IL-8 levels were measured by ELISA. **(L)** The interaction between rMICs and TLRs was evaluated by western blot. HEK293T cells transiently expressing TLR2-FLAG and TLR4-FLAG were lysed and incubated with His-rMIC1 (rMIC1^His^) or His-rMIC4 (rMIC4^His^). His-rMICs were subjected to Ni^2+^-affinity resin pull-down (lanes 6 to 9) and analysed for TLR2 and TLR4 binding by protein blotting with antibodies specific for FLAG-tag and then for rMIC (IgY, polyclonal). For these assays, rMIC1 and rMIC4 were passed through polymyxin B column, followed by incubation with polymyxin B sulphate salt media preparation that was added to the culture (see Material and Methods). Data in **(A-K)** are expressed as mean ±S.D. of triplicate wells and significance was calculated with ANOVA followed by Bonferroni’s multiple comparisons test. *p<0.05. Data are representative of three **(A-K)** and two **(L)** independent experiments.

### Cell activation induced by rMIC1 and rMIC4 results from the interaction of their CRDs with TLR2 and TLR4 N-glycans

We hypothesized that in order to trigger cell activation, rMIC1 and rMIC4 CRDs target oligosaccharides of the ectodomains of TLR2 (four N-linked glycans) [25] and TLR4 (nine N-linked glycans) [30]. This hypothesis was tested by stimulating BMDCs (Fig 4A) and BMDMs (Fig 4B) from WT mice with intact rMIC1 and rMIC4 or with the mutated forms of these microneme proteins, namely rMIC1-T126A/T220A and rMIC4-K469M, which lack carbohydrate binding activity [11, 18, 27]. IL-12 levels in culture supernatants were lower upon stimulation with rMIC1-T126A/T220A or rMIC4-K469M, showing that WT induction of cell activation requires intact rMIC1 and rMIC4 CRDs. The same microneme proteins were used to stimulate TLR2-transfected HEK293T cells (Fig 4C), and similarly, lower IL-8 production was obtained in response to mutated rMIC1 or rMIC4 compared to that seen in response to intact proteins. These observations demonstrated that rMIC1 and rMIC4 CRDs are also necessary for inducing HEK cell activation.

**Fig 4.**
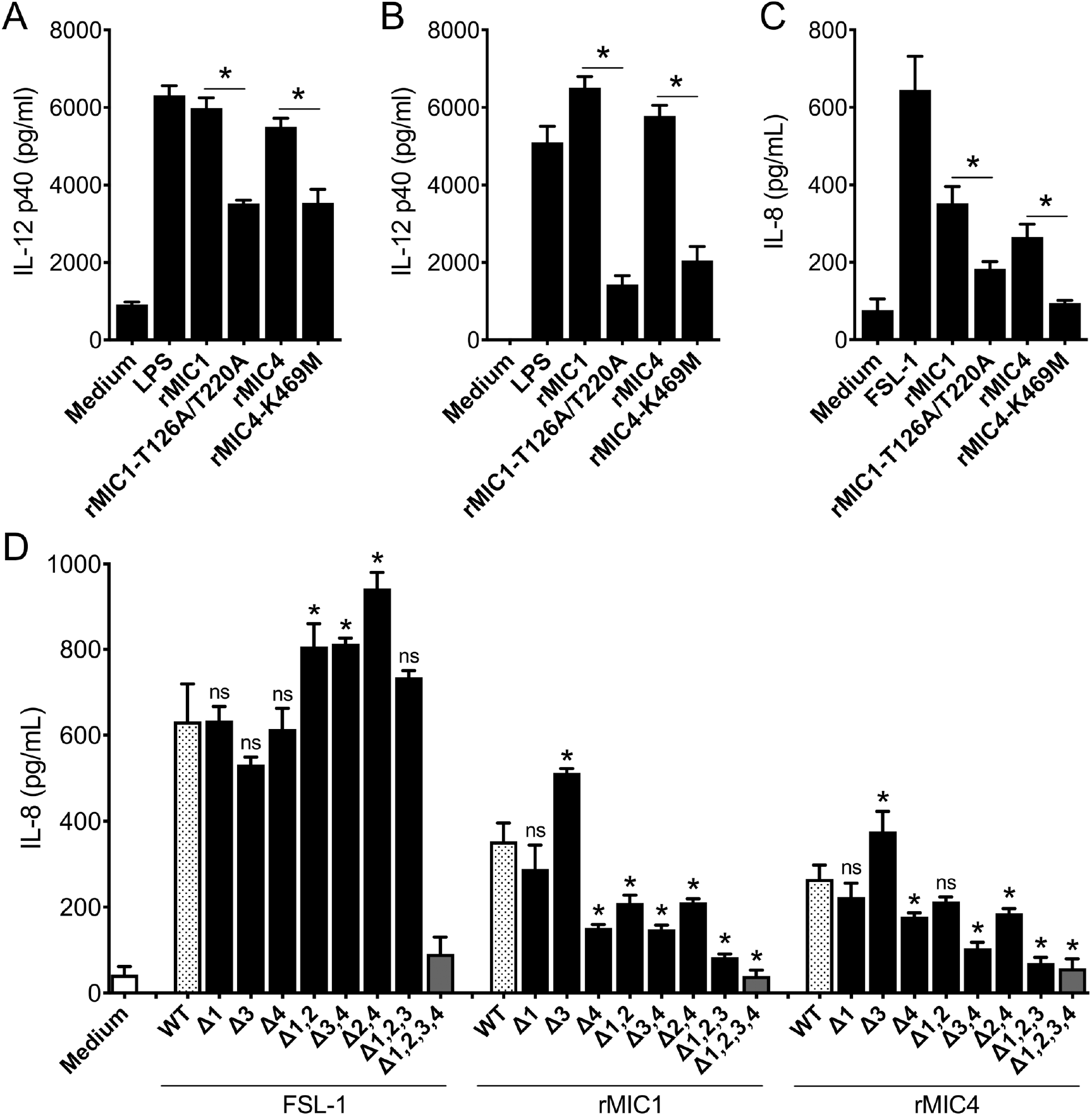
The cellular activation induced by rMICs via TLRs depends on carbohydrate recognition. **(A)** Bone marrow-derived macrophages and **(B)** bone marrow-derived dendritic cells from C57BL/6 mice and **(C)** transfected HEK293T cells expressing fully glycosylated TLR2 were stimulated with rMIC1 (WT) and rMIC4 (WT) or with their mutated forms, rMIC1-T126A/T220A and rMIC4-K469M, 5 μg/mL of each, for 48 h. LPS (100 ng/mL) and FSL-1 (100 ng/mL) were used as positive controls. IL-12p40 and IL-8 levels were measured by ELISA. **(D)** HEK293T cells expressing fully glycosylated TLR2 (with 4 N-glycans, WT) or glycosylation mutants of TLR2 (∆-1; ∆-4; ∆-1,2; ∆-3,4; ∆-2,4; ∆-1,2,3; ∆-1,2,3,4) were stimulated with rMIC1 or rMIC4. FSL-1 (100 ng/mL) was used as positive control. IL-8 levels were measured by ELISA. The statistical analysis compared fully glycosylated TLR2 (WT) and TLR2 mutants for the N-glycosylation sites for the same stimuli. For these assays, rMIC1, rMIC1-T126A/T220A, rMIC4 and rMIC4-K469M were passed through polymyxin B column, followed by incubation with polymyxin B sulphate salt media preparation that was added to the culture (see Material and Methods). Data are expressed as mean ±S.D. of triplicate wells and significance was calculated with ANOVA followed by Bonferroni’s multiple comparisons test. *p<0.05. Data are representative of three independent experiments.

We used an additional strategy to examine the ability of rMIC1 and rMIC4 to bind to TLR2 N-glycans. In this approach, HEK cells transfected with the fully N-glycosylated TLR2 ectodomain or with the TLR2 glycomutants [25] were stimulated with a control agonist (FSL-1) or with rMIC1 or rMIC4. HEK cells transfected with any TLR2 form, except those expressing totally unglycosylated TLR2 (mutant ∆1,2,3,4), were able to respond to FSL-1 (Fig 4D), a finding that is consistent with the previous report that the ∆1,2,3,4 mutant is not secreted by HEK293T cells [25]. Cells transfected with TLR2 lacking only the first or the third N-glycan (mutant ∆1; ∆3) responded to all stimuli. The response to the rMIC1 stimulus was significantly reduced in cells transfected with five different TLR2 mutants, lacking some combination of the second, third, and fourth N-glycans (Fig 4D). Moreover, rMIC4 stimulated IL-8 production was significantly reduced in cells transfected with the mutants lacking some combination of the third and fourth N-glycans (Fig 4D).

These results indicate that *T. gondii* MIC1 and MIC4 use their CRDs to induce TLR2- and TLR4-mediated cell activation. Among the TLR2 N-glycans, the rMIC1 CRD likely targets the second, third, and fourth glycan, whereas the rMIC4 CRD targets only the third and fourth. Additionally, our findings suggested that TLR2 and TLR4 activation is required to enhance the production of IL-12 by APCs following rMIC stimulation.

### The IL-12 production during *T. gondii in vitro* infection depends partially on MIC1 and MIC4 proteins and their ability to recognize carbohydrates on APCs surface

Because IL-12 production is induced by rMICs that engage TLR2 and TLR4 N-glycans expressed on innate immune cells, we investigated whether such production is impaired when APCs are infected with *T. gondii* lacking MIC1 and/or MIC4 proteins, as well as complemented strains expressing mutant versions of these proteins that fail to bind TLR2 or TLR4 carbohydrates. We generated Δ*mic1* and Δ*mic4* strains in an RH strain expressing GFP and Luciferase using CRISPR/Cas9 to replace the endogenous MIC gene with the drug-selectable marker HPT (HXGPRT – hypoxanthine-xanthine-guanine phosphoribosyl transferase) (Fig 5A and 5B). We then complemented MIC deficient parasites with mutated versions expressing an HA-tag, thus generating the Δ*mic1*::MIC1-T126A/T220A^HA^ or Δ*mic4*::MIC4-K469M^HA^ strains (Fig 5A) that expressed endogenous levels of MIC1 and MIC4 as confirmed by Western Blotting (Fig 5C).

**Fig 5.**
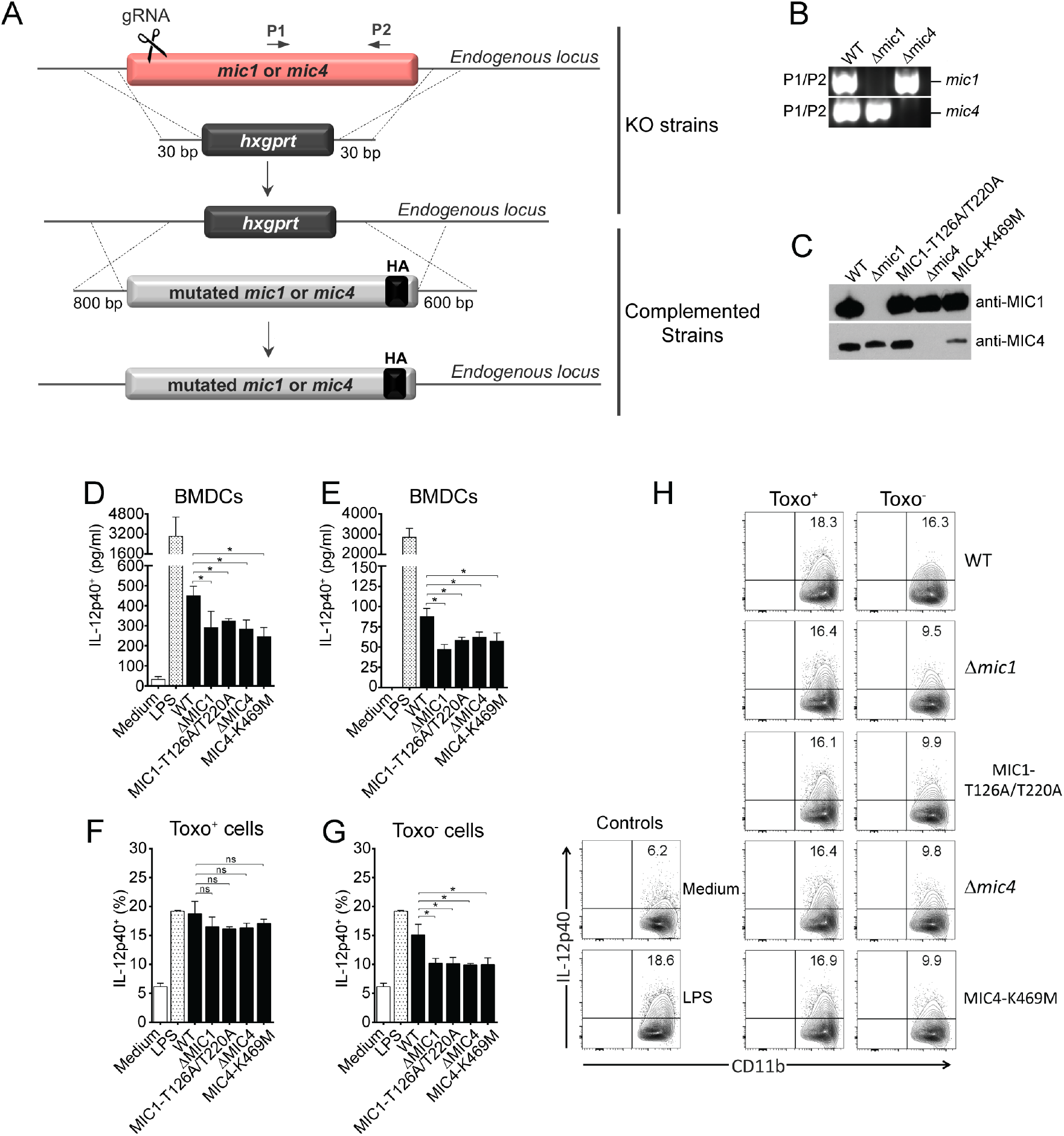
The IL-12 production during *T. gondii in vitro* infection partially depends on MICs and their ability to recognize carbohydrates on APCs surface. **(A)** Schematic representation of knockout and complementation constructs for MIC1 and MIC4 loci. The endogenous loci were disrupted using the hypoxanthine-xanthine-guanine phosphoribosyl transferase (HPT)-selectable marker and CRISPR methodology. **(B)** PCR analysis for MIC1 and MIC4 loci of gDNA from parental (WT RH-Δ*ku80*/Δ*hpt*-GFP/Luc) and knockout (RH-Δ*ku80*/Δ*mic1*-GFP/Luc and RH-Δ*ku80*/Δ*mic4*-GFP/Luc) strains. **(C)** Western blot analysis of an equal loading of whole cell lysates corresponding to 3 × 10^6^ tachyzoites (1 × 10^8^/mL) from WT, Δ*mic1*, Δ*mic1*::MIC1-T126A/T220A, Δ*mic4* and Δ*mic4*::K469M parasites. The membrane was probed with anti-MIC1 (IgY, 1:20,000) and anti-MIC4 (IgY, 1:8,000). **(D)** Bone marrow-derived dendritic cells (BMDCs) and **(E)** Bone marrow-derived macrophages (BMDMs) from C57BL/6 were infected with WT, Δ*mic1*, Δ*mic1*::MIC1-T126A/T220A, Δ*mic4* and Δ*mic4*::K469M strains (MOI 3). LPS (100 ng/mL) was used as positive control. Cell-culture supernatants were collected 24 hours post-infection and the IL-12p40 production was analyzed by ELISA. **(F and G)** Frequency of IL-12p40^+^ BMDCs (CD11b^+^IL-12p40^+^) after 20-24 hours of *in vitro* infection with WT, Δ*mic1*, Δ*mic1*::MIC1-T126A/T220A, Δ*mic4* and Δ*mic4*::K469M strains (MOI 1). Brefeldin A was added to the culture for 8 hours. LPS (100 ng/mL) was used as positive control. **(H)** Representative dot plots showing IL-12p40 staining in *T. gondii* infected or non-infected (SAG1^+^ or SAG1^−^) CD11b^+^ cells after 20-24 hours of *in vitro* infection with WT, Δ*mic1*, Δ*mic1*::MIC1-T126A/T220A, Δ*mic4* and Δ*mic4*::K469M strains (MOI 1). Brefeldin A was added to the culture for 8 hours. LPS (100 ng/mL) was used as positive control. Data are expressed as mean ±S.D. of triplicate wells and significance was calculated with ANOVA followed by Bonferroni’s multiple comparisons test. *p<0.05. Data are representative of three **(D and E)** and two **(F-H)** independent experiments.

IL-12 secretion by BMDCs and BMDMs infected with WT, Δ*mic1*, Δ*mic1*::MIC1-T126A/T220A, Δ*mic4* and Δ*mic4*::K469M parasites was assessed at 24 hours post infection. All mutant strains (Δ*mic1*, Δ*mic1*::MIC1-T126A/T220A, Δ*mic4* and Δ*mic4*::K469M) induced lower IL-12 secretion by BMDCs (Fig 5D) and BMDMs (Fig 5E) compared to that induced by WT parasites, indicating that engagement of TLR2 and TLR4 cell surface receptors by the MIC lectin-specific activity led to an early release of IL-12.

Using flow cytometry, we confirmed that parasites deficient in MIC1or MIC4, or mutated in their carbohydrate recognition domain resulted in lower intracellular IL-12 production than WT infected BMDCs (Fig 5F-5H). Interestingly, the Toxo^+^ BMDCs presented the same level of intracellular IL-12, independent of the *T. gondii* strain infected (Fig 5F and 5H). Whereas the Toxo^−^ BMDCs produced less IL-12 when they were infected with knockout or CRD-mutated *T. gondii* compared to WT-infected cells (Fig 5G and 5H). Taken altogether, these results indicate that MIC1 and MIC4 induce IL-12 production in innate immune cells during *in vitro T. gondii* infection. It is known that other parasite factors act as IL-12 inducers, such as profilin, which is a TLR11 and TLR12 agonist [29, 31], or GRA7 [32], GRA15 [33], and GRA24 [34], which directly trigger intracellular signalling pathways in a TLR-independent manner, and these likely account for the majority of IL-12 released after 24 hours of intracellular infection.

### MIC1, but not MIC4, contributes to the cytokine storm and acute death during *in vivo* murine infection with *T. gondii*

Given the importance of MIC1 and MIC4 as lectins that engage TLR2 and TLR4 N-glycans to induce increased levels of IL-12 release during *T. gondii in vitro* infection, we investigated the biological relevance of these proteins during *in vivo* infection. Mice were injected with 50 tachyzoites of RH WT, Δ*mic1*, Δ*mic1*::MIC1-T126A/T220A, Δ*mic4* and Δ*mic4*::MIC4-K469M strains into the peritoneum of CD-1 outbred mice, a lethal dose that causes acute mortality. The survival curve showed that parasites deficient in MIC1 (Δ*mic1* group) or mutated to remove MIC1 lectin binding activity (Δ*mic1*::MIC1-T126A/T220A group) were less virulent, resulting in a slight, but significant (p=0.0017) increase in mouse survival (12 days post-infection) compared to WT infected mice that all succumbed to infection by day 10 (Fig 6A). This was not the result of a difference in parasite load, which was equivalent across all *T. gondii*-infected mice at Day 5 (Fig 6D and 6I). Whereas, the absence of the MIC4 gene or MIC4 lectin activity did not change the survival curve (Fig 6E) indicating that MIC4 is less relevant than MIC1 during *in vivo* infection.

**Fig 6.**
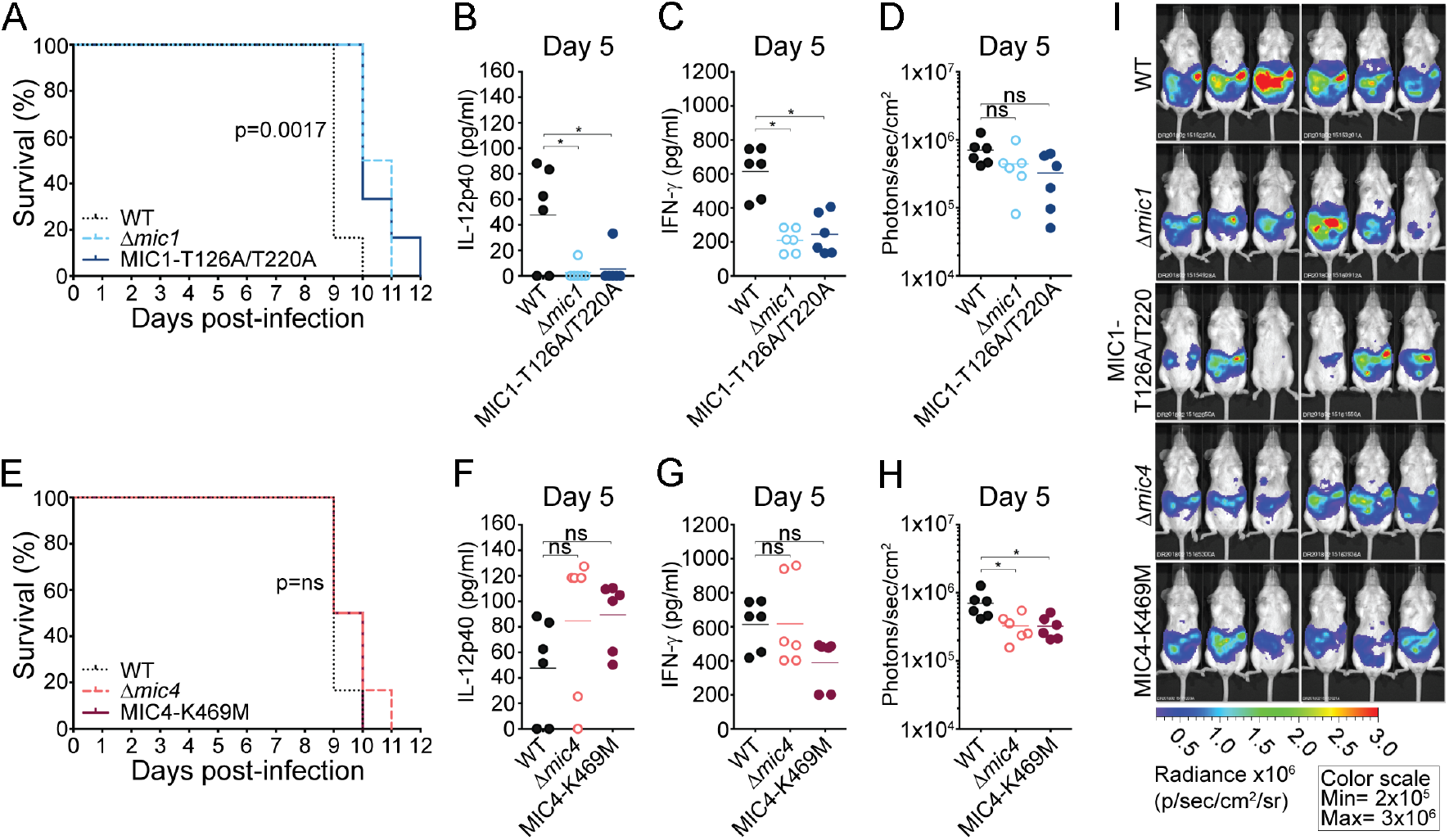
MIC1 lectin activity, but not MIC4, contributes to virulence in mice during *in vivo* infection with *T. gondii*. CD-1 mice were infected intraperitoneally with RH engineered strains of *T. gondii* at an infectious dose of 50 tachyzoites/mouse (n=6). Mortality kinetics of mice infected with **(A)** WT, Δ*mic1* and Δ*mic1*::MIC1-T126A/T220A strains or **(E)** WT, Δ*mic4* and Δ*mic4*::MIC4-K469M parasites. At day 5 post-infection the sera were collected for measuring systemic **(B and F)** IL-12p40 and **(C and G)** IFN-γ. **(D, H and I)** Bioluminescent detection in photons/sec/cm^2^ shows parasite burden 5 days post-infection. Data are expressed as mean ±S.D. and significance was calculated with ANOVA followed by Bonferroni’s multiple comparisons test. *p<0.05. Data are representative of three independent experiments, with total n=16.

Acute mortality in CD-1 mice infected with Type I *T. gondii* is related to the induction of a cytokine storm, mediated by high levels of IFN-γ production. Thus, we measured systemic levels of IFN-γ and IL-12 in mice infected with WT, Δ*mic1*, Δ*mic1*::MIC1-T126A/T220A, Δ*mic4* and Δ*mic4*::MIC4-K469M strains. According to Kugler et al. (2013), the peak of systemic IL-12p40 and IFN-γ during ME49-*T. gondii* infection is between days 5-6 post-infection, therefore, we measured these cytokines in the serum of CD-1-infected mice at day 5. Mice infected with Δ*mic1* or Δ*mic1*::MIC1-T126A/T220A strains had 3-5 fold lower systemic levels of IL-12 (Fig 6B; p=0.016) and IFN-γ (Fig 6C; p≤0.0002) than WT infected mice. In contrast, mice infected with parasites lacking the MIC4 gene, or those expressing the mutant version of MIC4 showed no difference in IL-12 (Fig 6F) or IFN-γ (Fig 6G) compared to WT infected mice. Hence, only MIC1 altered systemic levels of key cytokines induced during *T. gondii in vivo* infection, and mice survived longer with lower systemic levels of cytokines typically associated with acute mortality.

### MIC1 wild type complemented strain restores the cytokine storm and acute mortality kinetics during *in vivo* infection with *T. gondii*

To formally show that MIC1 alters systemic levels of pro-inflammatory cytokines associated with acute mortality, we complemented ∆*mic1* parasites at the endogenous locus with a Type I allele of MIC1 expressing an HA tag (MIC1^HA^). Western blotting for either MIC1 or HA expression showed WT levels of MIC1 expression in the complemented parasites ∆*mic1*::MIC1^HA^ (Fig 7A). The complemented strain restored WT virulence kinetics during *in vivo* infection and all mice died acutely, in contrast to Δ*mic1* or Δ*mic1*::MIC1-T126A/T220A parasites, that had a slight, but significant delay in their acute mortality kinetics (Fig 7B; p=0.0082). Systemic levels of IFN-γ (Fig 7C) and parasite load (Fig 7D and 7E) from mice infected with the complemented strain were indistinguishable from WT. To better resolve the apparent difference in acute mortality, parasites were injected into the right footpad to monitor mouse weight loss and survival kinetics [35]. Mice infected locally in the footpad with Δ*mic1* survived significantly longer, or did not die (Fig 7G; p=0.0031), and lost less weight during acute infection (Fig 7F) than those infected with WT or Δ*mic1*::MIC1 complemented parasites. Further, mice infected with Δ*mic1*::MIC1-T126A/T220A parasites that fail to bind TLR2 and TLR4 N-glycans *in vivo* also lost less weight and survived significantly longer than WT or Δ*mic1*::MIC1 complemented parasites (Fig 7F and G). In conclusion, our results suggest that MIC1 operates in two distinct ways; as an adhesin protein that promotes parasite infection competency, and as a lectin that engages TLR N-glycans to induce a stronger proinflammatory immune response, one that is unregulated and results in acute mortality upon RH infection of CD-1 mice.

**Fig 7.**
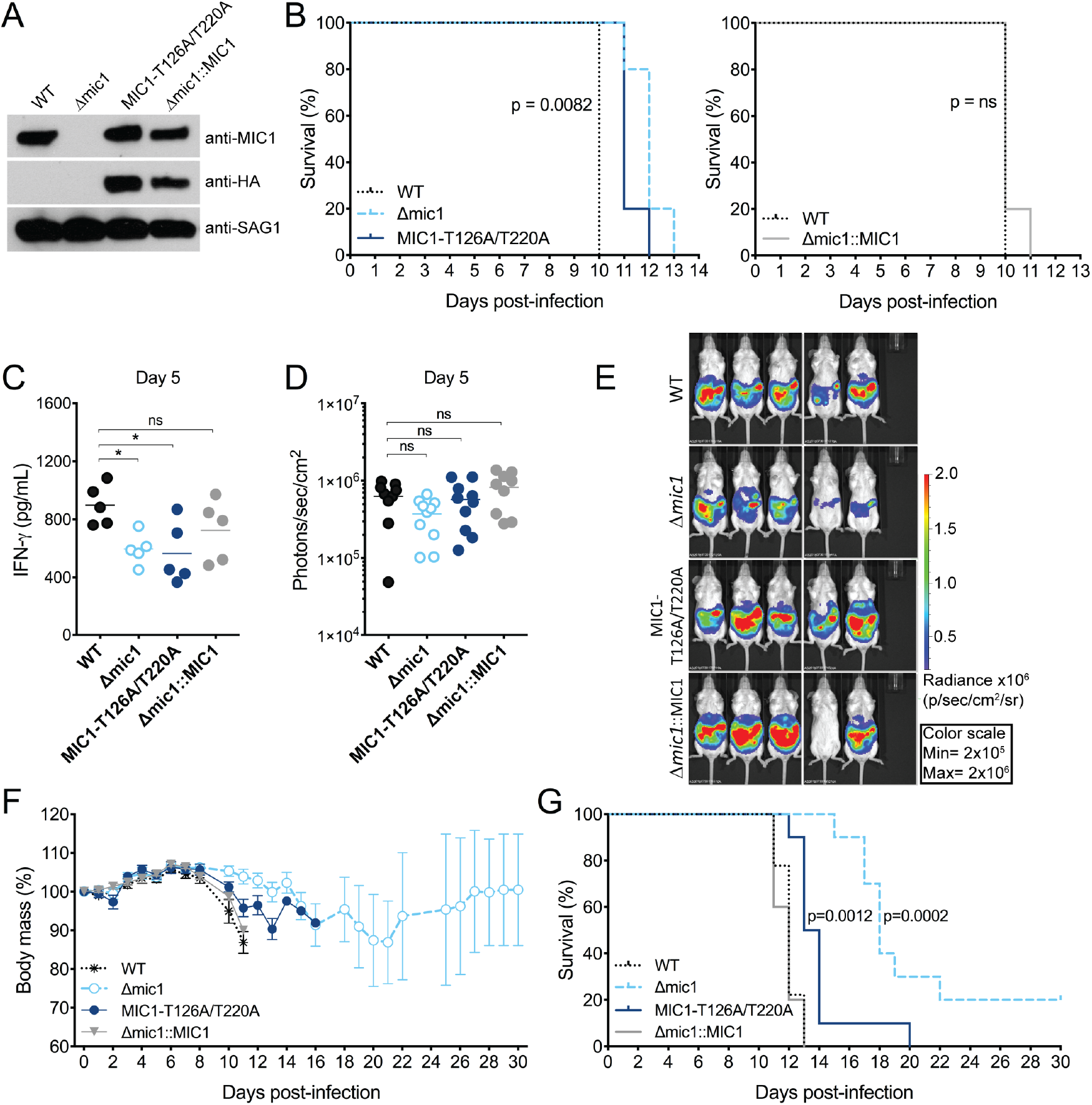
MIC1 wild type complemented strain restores virulence in mice during *in vivo* infection with *T. gondii*. **(A)** Western blot analysis of an equal loading of whole cell lysates corresponding to 3 × 10^6^ tachyzoites (1 × 10^8^/mL) from WT, Δ*mic1*, Δ*mic1*::MIC1-T126A/T220A and Δ*mic1*::MIC1 parasites. The membrane was probed with anti-MIC1 (IgY, 1:20,000), anti-HA (rabbit, 1:5,000) and anti-SAG1 (rabbit, 1:10,000). **(B)** Mortality kinetics of CD-1 mice infected intraperitoneally with WT, Δ*mic1*, Δ*mic1*::MIC1-T126A/T220A and Δ*mic1*::MIC1 at an infectious dose of 50 tachyzoites/mouse (n=5). **(C)** At day 5 post-infection the sera were collected for measuring systemic IFN-γ. **(D and E)** Bioluminescent detection in photons/sec/cm^2^ shows parasite burden 5 days post-infection. **(F and G)** Body mass and mortality kinetics of CD-1 mice infected subcutaneously with WT, Δ*mic1*, Δ*mic1*::MIC1-T126A/T220A and Δ*mic1*::MIC1 using an infectious dose of 10^4^ tachyzoites/mouse. Data are expressed as mean ±S.D. and significance were calculated with ANOVA followed by Bonferroni’s multiple comparisons test. *p<0.05. Data are representative of two independent experiments, total n=10.

## DISCUSSION

In this study, we report a new function for MIC1 and MIC4, two *T. gondii* microneme proteins involved in the host-parasite relationship. We show that rMIC1 and rMIC4, by interacting directly with N-glycans of TLR2 and TLR4, trigger a noncanonical carbohydrate recognition-dependent activation of innate immune cells. This results in IL-12 secretion and the production of IFN-γ, a pivotal cytokine that mediates parasite clearance and the development of a protective T cell response [19, 22], but in some cases, as seen during RH infection of CD-1 mice, promotes a dysregulated cytokine storm and acute mortality, as seen during RH infection of CD-1 mice [36]. This MIC-TLR activation event explains, at least in part, the resistance conferred by rMIC1 and rMIC4 administration against experimental toxoplasmosis [20, 21].

*T. gondii* tachyzoites express microneme proteins either on their surface or secrete them in their soluble form. These proteins may form complexes, such as those of MIC1, MIC4, and MIC6 (MIC1/4/6), in which MIC6 is a transmembrane protein that anchors the two soluble molecules MIC1 and MIC4 [8]. Genetic disruption of each one of these three genes does not interfere with parasite survival [8] nor its interaction with, and attachment to, host cells [10]; however, MIC1 has been shown to play a role in invasion and contributes to virulence in mice [10]. We previously isolated soluble MIC1/4, a lactose-binding complex from soluble *T. gondii* antigens (STAg) [17], and its lectin activity was confirmed by the ability of MIC1 to bind sialic acid [9] and MIC4 to β-galactose [18]. We also reported that MIC1/4 stimulates adherent splenic murine cells to produce IL-12 at levels as high as those induced by STAg [20]. Recently, it was also demonstrated that MIC1, MIC4 and MIC6 are capable of inducing IFN-γ production from memory T cells in mice chronically infected with *T. gondii* [37]. Our data herein shows that MIC1/4 binds to and activates TLRs via a novel lectin-carbohydrate interaction, rather than by its cognate receptor-ligand binding groove, establishing precisely how the interactions of microneme protein(s) with defined glycosylated receptor(s) expressed on the host cell surface are capable of altering innate priming of the immune system.

To formally demonstrate the MIC1/MIC4 binding to glycosylated TLR cell surface receptors we generated recombinant forms of MIC1 and MIC4, which retained their specific sialic acid- and β-galactose-binding properties as indicated by the results of their binding to fetuin and asialofetuin as well as the glycoarray assay. Both recombinant MIC1 and MIC4 triggered the production of proinflammatory and anti-inflammatory cytokines in DCs and macrophages via their specific recognition of TLR2 and TLR4 N-glycans, as well as by signaling through MyD88 and, partially, TRIF. Importantly, our results establish how binding of rMIC1 and rMIC4 to specific N-glycans present on TLR2 and TLR4 induces cell activation through this novel lectin-carbohydrate interaction. The ligands for MIC1 and MIC4, α2-3-sialyllactosamine and β1-3-or β1-4-galactosamine, respectively, are terminal N-glycan residues found on a wide-spectrum of mammalian cell surface-associated glycoconjugates. Thus, it is possible that additional lectin-carbohydrate interactions may exist between MIC1/4 and other cell surface receptors beyond TLR2 and TLR4. Such interactions likely evolved to facilitate adhesion and promote the infection competency of a wide-variety of host cells infected by *T. gondii*, further underscoring how these proteins exist as important virulence factors [10] beyond immune priming. However, it is the immunostimulatory capacity of rMIC1 and rMIC4 to target N-glycans on the ectodomains of TLR2 and TLR4 that likely rationalizes how these microneme proteins function as a double-edged sword during *T. gondii* infection. Mice infected by Type I strains die acutely due to a failure to regulate the cytokine storm induced by high levels of IL-12 and IFN-γ [38, 39]. In this study, *T. gondii* Type I strains engineered to be deficient in MIC1 or defective in binding TLR2/4 N-glycans lost less weight, survived significantly longer, and produced less IL-12 and IFN-γ. Future studies that test whether the immunostimulatory effect of MIC1/4 alters the pathogenesis and cyst burden of Type II strains of *T. gondii* should be pursued to formally demonstrate that Type II parasites rely on MIC1/4 induction of Th1-biased cytokines in order to limit tachyzoite proliferation and induce a life-long persistent bradyzoite infection.

Several pathogens are known to synthesize lectins, which are most frequently reported to interact with glycoconjugates on host cells to promote adherence, invasion, and colonization of tissues [40–43]. Nonetheless, there are currently only a few examples of lectins from pathogens that recognize sugar moieties present in TLRs and induce IL-12 production by innate immune cells. Paracoccin, a GlcNAc-binding lectin from the human pathogen *Paracoccidioides brasiliensis*, induces macrophage polarization towards the M1 phenotype [44] and the production of inflammatory cytokines through its interaction with TLR2 N-glycans [45]. Furthermore, the galactose-adherence lectin from *Entamoeba histolytica* activates TLR2 and induces IL-12 production [46]. In addition, the mammalian soluble lectin SP-A, found in lung alveoli, interacts with the TLR2 ectodomain [47]. The occurrence of cell activation and IL-12 production as a consequence of the recognition of TLR N-glycans has also been demonstrated using plant lectins with different sugar-binding specificities [48, 49].

The binding of MIC1 and MIC4, as well as the lectins above, to TLR2 and TLR4 may be associated with the position of the specific sugar residue present on the receptor’s N-glycan structure. Since the N-glycan structures of TLR2 and TLR4 are still unknown, we assume that the targeted MIC1 and MIC4 residues, e.g. sialic acid α2-3-linked to galactose β1-3- and β1-4-galactosamines, are appropriately placed in the receptors’ oligosaccharides to allow the recognition phenomenon and trigger the activation of innate immune responses.

Several *T. gondii* proteins have previously been shown to activate innate immune cells in a TLR-dependent manner, but independent of sugar recognition. This is the case for profilin (TgPRF), which is essential for the parasite’s gliding motility based on actin polymerization; it is recognized by TLR11 [29] and TLR12 [31, 50]. In addition, *T. gondii*-derived glycosylphosphatidylinositol anchors activate TLR2 and TLR4 [51], and parasite RNA and DNA are ligands for TLR7 and TLR9, respectively [19, 22, 50]. The stimulation of all of these TLRs culminate in MyD88 activation which results in IL-12 production [19, 22]. Several other *T. gondii* secreted effector proteins regulate the production of proinflammatory cytokines such as IL-12, independent of TLRs. For example, the dense granule protein 7 (GRA7) induces MyD88-dependent NF-kB activation, which facilitates IL-12, TNF-α, and IL-6 production [32]. MIC3 is reported to induce TNF-α secretion and macrophage M1 polarization [52], whereas GRA15 expressed by Type II strains activates NF-kB, promoting the release of IL-12 [33], and GRA24 triggers the autophosphorylation of p38 MAP kinase and proinflammatory cytokine and chemokine secretion [34]. In contrast, TgIST interferes with IFN-γ induction by actively inhibiting STAT1-dependent proinflammatory gene expression indicating that the parasite is capable of both activating as well as inhibiting effector arms of the host immune response to impact its pathogenesis *in vivo* [53]. Thus, multiple secretory effector proteins of *T. gondii*, including MIC1 and MIC4, appear to work in tandem to ultimately promote protective immunity by either inducing or dampening the production of proinflammatory cytokines, the timing of which is central to controlling both the parasite’s proliferation during the acute phase of infection and the induction of an effective immune response capable of establishing a chronic infection [19].

Our results regarding soluble MIC1 and MIC4 confirmed our hypothesis that these two effector proteins induce the innate immune response against *T. gondii* through TLR2- and TLR4-dependent pathways. This is consistent with previous studies that highlight the importance of TLR signaling, as well as the MyD88 adapter molecule, as essential for conferring resistance to *T. gondii* infection [29, 51, 54, 55]. In addition, we show that both MIC1 and MIC4 on the parasite surface contribute to the secretion of IL-12 by macrophages and DCs during *in vitro* infection, but only MIC1 plays a significant role during *in vivo* infection, demonstrated by its ability to promote a dysregulated induction of systemic levels of IFN-γ and a proinflammatory cytokine storm that leads to acute mortality during murine infection.

## METHODS

### Ethics statement

All experiments were conducted in accordance to the Brazilian Federal Law 11,794/2008 establishing procedures for the scientific use of animals, and State Law establishing the Animal Protection Code of the State of Sao Paulo. All efforts were made to minimize suffering, and the animal experiments were approved by the Ethics Committee on Animal Experimentation (*Comissão de Ética em Experimentação Animal* – CETEA) of the Ribeirao Preto Medical School, University of Sao Paulo (protocol number 065/2012), following the guidelines of the National Council for Control of Animal Experimentation (*Conselho Nacional de Controle de Experimentação Animal* – CONCEA).

### Lac^+^ fraction and recombinant MIC1 and MIC4

The lactose-eluted (Lac^+^) fraction was obtained as previously reported [17, 21]. Briefly, the total soluble tachyzoite antigen (STAg) fraction was loaded into a lactose column (Sigma-Aldrich, St. Louis, MO) and equilibrated with PBS containing 0.5 M NaCl. The material adsorbed to the resin was eluted with 0.1 M lactose in equilibrating buffer and dialyzed against ultrapure water. The obtained fraction was denoted as Lac^+^ and confirmed to contain MIC1 and MIC4. For the recombinant proteins, rMIC1 and rMIC4 sequences were amplified from cDNA of the *T. gondii* strain ME49 with a 6-histidine tag added on the N-terminal, cloned into pDEST17 vector (Gateway Cloning, Thermo Fisher Scientific Inc., Grand Island, NY), and used to transform DH5α *E*. *coli* chemically competent cells for ampicillin expression selection, as described before [21]. The plasmids with rMIC1-T126A/T220A and rMIC4-K469M were synthesized by GenScript (New Jersey, US) using a pET28a vector, and the MIC sequences carrying the mutations were cloned between the *Nde*I and *Bam*H I sites. All plasmids extracted from DH5α *E*. *coli* were transformed in *E*. *coli* BL21-DE3 chemically competent cells to produce recombinant proteins that were then purified from inclusion bodies and refolded by gradient dialysis, as described previously for rMIC1 and rMIC4 wild type forms [21]. Endotoxin concentrations were measured in all protein samples using the Limulus Amebocyte Lysate Kit – QCL-1000 (Lonza, Basel, Switzerland). The rMIC1, rMIC1-T126A/T220A, rMIC4 and rMIC4-K469M contained 7.2, 3.2, 3.5 and 1.1 EU endotoxin/µg of protein, respectively. Endotoxin was removed by passing over two polymyxin-B columns (Affi-Prep Polymyxin Resin; Bio-Rad, Hercules, CA). Additionally, prior to all *in vitro* cell-stimulation assays, the proteins samples were incubated with 50 μg/mL of polymyxin B sulphate salt (Sigma-Aldrich, St. Louis, MO) for 30 min at 37 °C to remove possible residual LPS.

### Glycan array

The carbohydrate-binding profile of microneme proteins was determined by Core H (Consortium for Functional Glycomics, Emory University, Atlanta, GA), using a printed glycan microarray, as described previously [56]. Briefly, rMIC1-Fc, rMIC4-Fc, and Lac^+^-Fc in binding buffer (1% BSA, 150 mM NaCl, 2 mM CaCl_2_, 2 mM MgCl_2_, 0.05% (w/v) Tween 20, and 20 mM Tris-HCl, pH 7.4) were applied onto a covalently printed glycan array and incubated for 1 hour at 25 °C, followed by incubation with Alexa Fluor 488-conjugate (Invitrogen, Thermo Fisher Scientific Inc., Grand Island, NY). Slides were scanned, and the average signal intensity was calculated. The common features of glycans with stronger binding are depicted in Fig. 1a. The average signal intensity detected for all of the glycans was calculated and set as the baseline.

### Sugar-inhibition assay

Ninety-six-well microplates were coated with 1 μg/well of fetuin or asialofetuin, glycoproteins diluted in 50 μL of carbonate buffer (pH 9.6) per well, followed by overnight incubation at 4 °C. Recombinant MIC1 or MIC4 proteins (both wild type (WT) and mutated forms), previously incubated or not with their corresponding sugars, i.e. α(2-3)-sialyllactose for MIC1 and lacto-N-biose for MIC4 (V-lab, Dextra, LA, UK), were added into coated wells and incubated for 2 h at 25 °C. After washing with PBS, *T. gondii*-infected mouse serum (1:50) was used as the source of the primary antibody. The assay was then developed with anti-mouse peroxidase-conjugated secondary antibody, and the absorbance was measured at 450 nm in a microplate-scanning spectrophotometer (Power Wave-X; BioTek Instruments, Inc., Winooski, VT).

### Mice and parasites

Female C57BL/6 (WT), MyD88^−/−^, TRIF^−/−^, TLR2^−/−^, TLR3^−/−^, TLR4^−/−^, double knockout (DKO) TLR2^−/−^/TLR4^−/−^, TLR5^−/−^, and TLR9^−/−^ mice (all from the C57BL/6 background), 8 to 12 weeks of age, were acquired from the University of São Paulo – Ribeirão Preto campus animal facility, Ribeirão Preto, São Paulo, Brazil, and housed in the animal facility of the Department of Cell and Molecular Biology – Ribeirão Preto Medical School, under specific pathogen-free conditions. The TLR11^−/−^/TLR12^−/−^ DKO mice were maintained at American Association of Laboratory Animal Care-accredited animal facilities at NIAID/NIH. For the *in vivo* infections, female CD-1 outbred mice, 6 weeks of age were acquired from Charles River Laboratories, Germantown, MD, USA.

A clonal isolate of the *T. gondii* RH-Δ*ku80*/Δ*hpt* strain was used to generate the GFP/Luciferase strain, which was the recipient strain to generate the single-knockout parasites. The GFP/Luc sequence was inserted into the UPRT locus of *Toxoplasma* by double crossover homologous recombination using CRISPR/Cas-based genome editing and selected for FUDR resistance to facilitate the targeted GFP/Luc gene cassette knock-in. The MIC1 and MIC4 genes were replaced by the drug-selectable marker *hpt* (*hxgprt* – hypoxanthine-xanthine-guanine phosphoribosyl transferase) flanked by LoxP sites. For all gene deletions, 30 μg of guide RNA was transfected along with 15 µg of a repair oligo. Parasites were transfected and selected as previously described [57, 58]. For the MIC gene complementation, the sequence was amplified from RH genomic DNA with the addition of one copy of HA-tag sequence (TACCCATACGATGTTCCAGATTACGCT) before the stop codon, and cloned into pCR2.1-TOPO vector, followed by site-directed mutagenesis using the Q-5 kit (New England Biolabs) in order to generate point mutations into MIC1 (MIC1-T126A/T220A) and MIC4 (MIC4-K469M) sequences. For transfections, 30 μg of guide RNA was transfected along with 20 µg of linearized pTOPO vector containing the MIC mutated sequences.

Strains were maintained in human foreskin fibroblast (HFF) cells grown in Dulbecco’s modified Eagle’s medium (DMEM) supplemented with 10% heat-inactivated foetal bovine serum (FBS), 0.25 mM gentamicin, 10 U/mL penicillin, and 10 μg/mL streptomycin (Gibco, Thermo Fisher Scientific Inc., Grand Island, NY).

### Bone marrow-derived dendritic cells and macrophages

Bone marrows of WT, MyD88^−/−^, TRIF^−/−^, TLR2^−/−^, TLR3^−/−^, TLR4^−/−^, DKO TLR2^−/−^/TLR4^−/−^, TLR5^−/−^, TLR9^−/−^, and DKO TLR11^−/−^/TLR12^−/−^ mice were harvested from femurs and hind leg bones. Cells were washed with RPMI medium and resuspended in RPMI medium with 10% FBS, 10 U/mL penicillin, and 10 μg/mL streptomycin (Gibco). For dendritic cell (DC) differentiation, we added 10 ng/mL of recombinant murine GM-CSF (Prepotech, Rocky Hill, NJ), and 10 ng/mL murine recombinant IL-4 (eBioscience, San Diego, CA); for macrophage differentiation, 30% of L929 conditioned medium was added to RPMI medium with 10% FBS. The cells were cultured in 100 × 20 mm dish plates (Costar; Corning Inc., Corning, NY), supplemented with respective conditioned media at days 3 and 6 for DCs, and at day 4 for macrophages. DCs were incubated for 8–9 days and macrophages for 7 days; the cells were then harvested and plated into 24-well plates at 5 × 10^5^ cells/well for protein stimulations or *T. gondii* infections, followed by ELISA. Cell purity was analyzed by flow cytometry. Eighty-five percent of differentiated dentritic cells were CD11b^+^/CD11c^+^, while 94% of differentiated macrophages were CD11b^+^.

### HEK293T cells transfection

Human embryonic kidney 293T (HEK293T) cells, originally acquired from American Tissue Culture Collection (ATCC, Rockville, MD), were used as an expression tool [59] for TLR2 and TLR4 [45, 60]. The cells grown in DMEM supplemented with 10% FBS (Gibco), and were seeded at 3.5 × 10^5^ cells/mL in 96-well plates (3.5 × 10^4^ cells/well) 24 h before transfection. Then, HEK293T cells were transiently transfected (70-80% confluence) with human TLR2 plasmids as described previously [25] or with CD14, CD36, MD-2 and TLR4 [61] using Lipofectamine 2000 (Invitrogen) with 60 ng of NF-κB Luc, an NF-κB reporter plasmid, and 0.5 ng of *Renilla* luciferase plasmid, together with 60 ng of each gene of single and multiple glycosylation mutants and of TLR2 WT genes [25]. After 24 h of transfection, the cells were stimulated overnight with positive controls: P3C (Pam3CSK4; EMC Microcollections, Tübingen, Germany), fibroblast stimulating ligand-1 (FSL-1; EMC Microcollections), or LPS Ultrapure (standard LPS, *E. coli* 0111:B4; Sigma-Aldrich); or with the negative control for cell stimulation (the medium). Cells transfected with empty vectors, incubated either with the medium or with agonists (FSL-1 or P3C), were also assayed; negative results were required for each system included in the study. IL-8 was detected in the culture supernatants. The absence of Mycoplasma contamination in the cell culture was certified by indirect fluorescence staining as described previously [62].

### Cytokine measurement

The quantification of human IL-8 and mouse IL-12p40, IL-6, TNF-α, and IL-10 in the supernatant of the cultures was performed by ELISA, following the manufacturer’s instructions (OptEIA set; BD Biosciences, San Jose, CA). Human and murine recombinant cytokines were used to generate standard curves and determine cytokine concentrations. The absorbance was read at 450 nm using the Power Wave-X spectrophotometer (BioTek Instruments).

### TLR2-FLAG and TLR4-FLAG plasmids

The pcDNA4/TO-FLAG plasmid was kindly provided by Dr. Dario Simões Zamboni. The pcDNA4-FLAG-TLR2 and pcDNA4-FLAG-TLR4 plasmids were constructed as follows. RNA from a P388D1 cell line (ATCC, Rockville, MD) was extracted and converted to cDNA with Maxima H Minus Reverse Transcriptase (Thermo-Fisher Scientific, Waltham, MA USA) and oligo(dT). TLR2 and TLR4 were amplified from total cDNA from murine macrophages by using Phusion High-Fidelity DNA Polymerase and the phosphorylated primers TLR2_F: ATGCTACGAGCTCTTTGGCTCTTCTGG, TLR2_R: CTAGGACTTTATTGCAGTTCTCAGATTTACCCAAAAC, TLR4_F: TGCTTAGGATCCATGATGCCTCCCTGGCTCCTG and TLR4_R: TGCTTAGCGGCCGCTCAGGTCCAAGTTGCCGTTTCTTG. The fragments were isolated from 1% agarose/Tris-acetate-ethylenediaminetetraacetic acid gel, purified with GeneJET Gel Extraction Kit (Thermo-Fisher Scientific), and inserted into the pcDNA4/TO-FLAG vector by using the restriction enzymes sites for NotI and XbaI (Thermo-Fisher Scientific) for TLR2, and BamHI and NotI (Thermo-Fisher Scientific) for TLR4. Ligation reactions were performed by using a 3:1 insert/vector ratio with T4 DNA Ligase (Thermo-Fisher Scientific) and transformed into chemically competent *Escherichia coli* DH5α cells. Proper transformants were isolated from LB agar medium plates under ampicillin selection (100 μg/mL) and analyzed by PCR, restriction fragment analysis, and DNA sequencing. All reactions were performed according to the manufacturer’s instructions.

### Pull-down assay and Western Blot

We used the lysate of HEK293T cells transfected (70-80% confluence) with plasmids containing TLR2-FLAG or TLR4-FLAG. After 24 h of transfection, the HEK cells were lysed with a non-denaturing lysis buffer (20 mM Tris, pH 8.0, 137 mM NaCl, and 2 mM EDTA) supplemented with a protease inhibitor (Roche, Basel, Switzerland). After 10 min of incubation on ice, the lysate was subjected to centrifugation (16,000 *g*, at 4 °C for 5 min). The protein content in the supernatant was quantified by the BCA method, aliquoted, and stored at −80 °C. For the pull-down assay, 100 μg of the lysate from TLR2-FLAG- or TLR4-FLAG-transfected HEK cells were incubated with 10 μg of TgMIC1 or TgMIC4 overnight at 4 °C. Since these proteins had a histidine tag, the samples were purified on nickel-affinity resin (Ni Sepharose High Performance; GE Healthcare, Little Chalfont, UK) after incubation for 30 min at 25 °C and centrifugation of the fraction bound to nickel to pull down the TgMIC-His that physically interacted with TLR-FLAG (16,000 *g*, 4 °C, 5 min). After washing with PBS, the samples were resuspended in 100 μL of SDS loading dye with 5 μL of 2-mercaptoethanol, heated for 5 min at 95 ºC, and 25 μL of total volume was run on 10% SDS-PAGE. After transferring to a nitrocellulose membrane (Millipore, Billerica, MA), immunoblotting was performed by following the manufacturer’s protocol. First, the membrane was incubated with anti-FLAG monoclonal antibodies (1:2,000) (Clone G10, ab45766, Sigma-Aldrich) to detect the presence of TLR2 or TLR4. The same membrane was then subjected to secondary probing and was developed with anti-TgMIC1 (IgY; 1:20,000) or anti-TgMIC4 (IgY; 1:8,000) polyclonal antibodies and followed by incubation with secondary polyclonal anti-chicken IgY-HRP (1:4,000) (A9046, Sigma-Aldrich) to confirm the presence of TgMIC1 and TgMIC4.

### *In vitro* infections

Bone marrow-derived dendritic cells (BMDCs) and bone marrow-derived macrophages (BMDMs) were infected with WT (Δ*ku80*/Δ*hpt*), Δ*mic1*, Δ*mic1*::MIC1-T126A/T220A, Δ*mic4* or Δ*mic4*::MIC4-K469M (Type I, RH background) strains recovered from T25 flasks with HFF cell cultures. The T25 flasks were washed with RPMI medium to completely remove parasites, and the collected material was centrifuged for 5 min at 50 *g* to remove HFF cell debris. The resulting pellet was discarded, and the supernatant containing the parasites was centrifuged for 10 min at 1,000 *g* and resuspended in RPMI medium for counting and concentration adjustments. BMDCs and BMDMs were dispensed in 24-well plates at 5 × 10^5^ cells/well (in RPMI medium supplemented with 10% FBS), followed by infection with 3 parasites per cell (multiplicity of infection, MOI 3). Then, the plate was centrifuged for 3 min at 200 *g* to synchronize the contact between cells and parasites and incubated at 37 °C. The supernatants were collected at 6, 12, 24, and 48 h after infection for quantification of IL-12p40.

### *In vivo* infections and Luciferase assay

Six-week-old female CD-1 outbred mice were infected by intraperitoneal injection with 50 tachyzoites of RH engineered strains diluted in 500 µl of phosphate-buffered saline. The mice were weighed daily and survival was evaluated Bioluminescent detection of firefly luciferase activity was performed at day 5 post-infection using an IVIS BLI system from Xenogen to monitor parasite burden. Mice were injected with 3 milligrams (200 µl) of D-luciferin (PerkinElmer) substrate, and after 5 minutes the mice were imaged for 300 seconds to detect the photons emitted.

### Statistical analysis

The data were plotted and analysed using GraphPad Prism 7.0 software (GraphPad, La Jolla, CA). Statistical significance of the obtained results was calculated using analysis of variance (One-way ANOVA) followed by Bonferroni’s multiple comparisons test. Differences were considered significant when the *P* value was <0.05.

## ACKNOWLEDGEMENTS

We are grateful to Dr. Tiago Wilson Patriarca Mineo (Universidade Federal de Uberlândia – MG) for kindly provided us the wild type *Toxoplasma gondii* (RH); to Dr. Larissa Dias Cunha and Dr. Dario Simões Zamboni (Universidade de São Paulo – SP) for help with double knockout TLR2/TLR4 mice generation and for kindly provide the pcDNA4/TO-FLAG plasmid; to Izaíra Tincani Brandão and Ana Paula Masson for technical assistance with endotoxin measurements; to Patricia Vendrusculo, Sandra Thomaz and Sara Hieny for all technical support essential for this study. We wish to acknowledge the grants from Consortium for Functional Glycomics (#GM62116), for doing the glycoarray assays; UK Medical Research Council (#G1000133 to N.J.G.), and Wellcome Investigator Award (#WT100321/z/12/Z to N.J.G.). This work was supported in part by the National Institute of Allergy and Infectious Diseases, National Institutes of Health (A.S.S. and M.E.G.).

## SUPPLEMENTARY INFORMATION

**S1 Fig.**
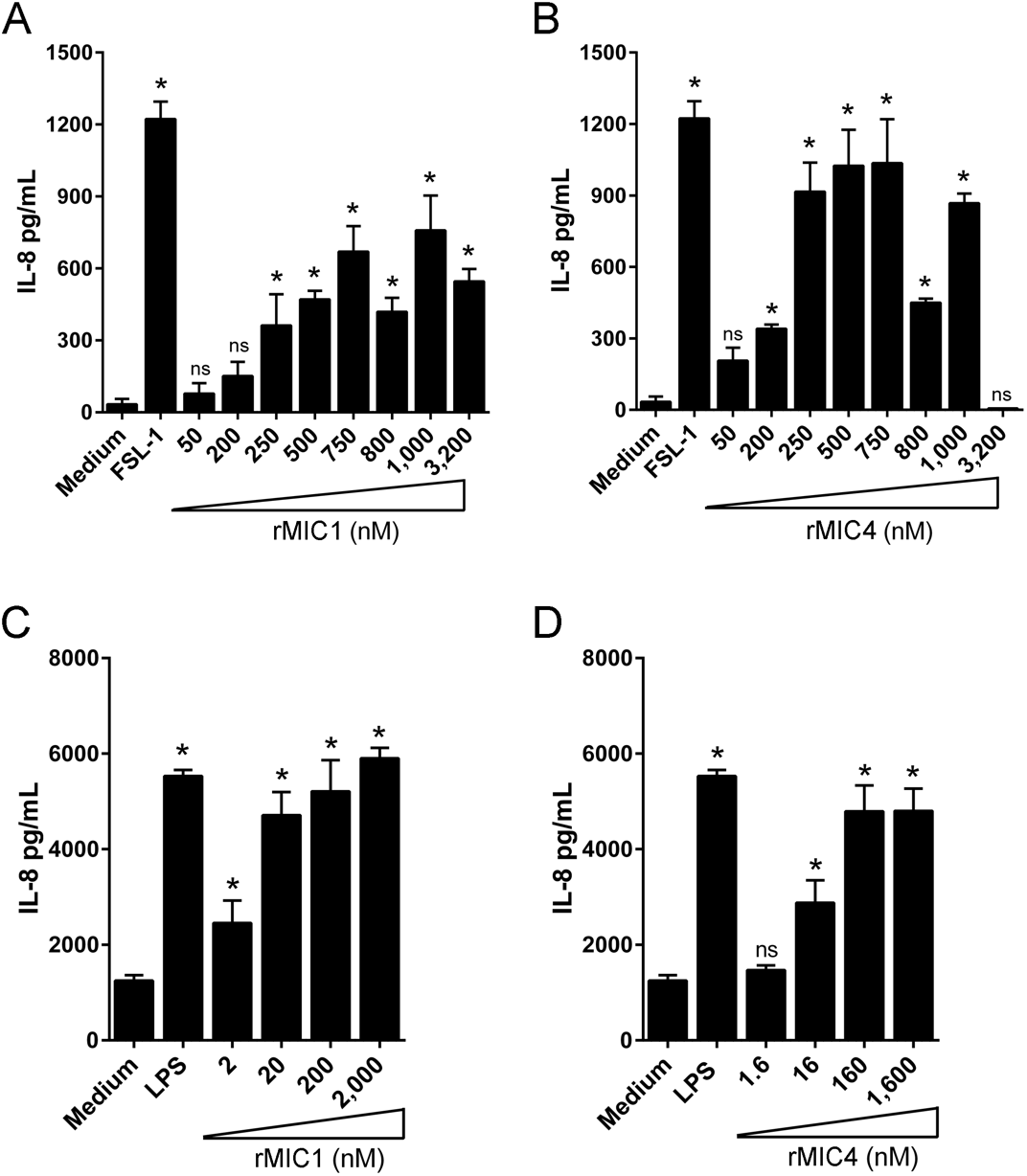
Effect of different concentrations of rMIC1 and rMIC4 on the transfected HEK cells. HEK293T cells expressing **(A and B)** TLR2 or **(C and D)** TLR4 were stimulated with increasing concentrations of **(A and C)** rMIC1 and **(B and D)** rMIC4 for 24 h. FSL-1 (100 ng/mL) LPS (100 ng/mL) were used as positive controls. IL-8 levels were measured by ELISA. Data are expressed as mean ±S.D. of triplicate wells and significance was calculated with ANOVA. *p<0.05. Data are representative of two independent experiments.

